# GABA-receptors are a new druggable target for limiting disease severity, lung viral load, and death in SARS-CoV-2 infected mice

**DOI:** 10.1101/2022.06.07.494579

**Authors:** Jide Tian, Barbara J. Dillion, Jill Henley, Lucio Comai, Daniel L. Kaufman

## Abstract

GABA-receptors (GABA-Rs) are well-known neurotransmitter receptors in the central nervous system. GABA-Rs are also expressed by immune cells and lung epithelial cells and GABA-R agonists/potentiators reduce inflammatory immune cell activities and limit acute lung injuries. Notably, plasma GABA levels are reduced in hospitalized COVID-19 patients. Hence, GABA-R agonists may have therapeutic potential for treating COVID-19. Here, we show that oral GABA treatment initiated just after SARS-CoV-2 infection, or 2 days later near the peak of lung viral load, reduced disease severity, lung coefficient index, and death rates in K18-hACE2 mice. GABA-treated mice had a reduced viral load in their lungs and displayed shifts in their serum cytokine and chemokine levels that are associated with better outcomes in COVID-19 patients. Thus, GABA-R activation had multiple beneficial effects in this mouse model which are also desirable for the treatment of COVID-19. A number of GABA-R agonists are safe for human use and can be readily tested in clinical trials with COVID-19 patients. We also discuss their potential for limiting COVID-19-associated neuroinflammation.

## Introduction

Despite the great success of vaccines to reduce serious illness due to COVID-19, this approach has limitations due to break-through infections, vaccine hesitancy, viral variants, and novel coronaviruses. Antiviral drugs can greatly help reduce the risk for severe disease and mortality due to COVID-19, however, these drugs may not become readily available in developing countries and they may be less effective against coronaviruses that emerge in the future. The identification of additional therapeutics that have established safety records, are inexpensive, and do not have special storage requirements could be especially helpful for reducing COVID-19-associated morbidity and mortality worldwide.

γ-aminobutyric acid type A receptors (GABA_A_-Rs) are a family of ligand-gated chloride channels which play key roles in neurodevelopment and neurotransmission in the central nervous system (CNS) (1-3). GABA_A_-Rs are also expressed by cells of the human and murine immune systems (e.g., (4-7)). The activation of GABA_A_-Rs on innate immune system cells such as macrophages, dendritic, and NK cells reduces their inflammatory activities and shifts them toward anti-inflammatory phenotypes (7-15). Moreover, GABA_A_-R agonists inhibit the inflammatory activities of human and murine Th17 and Th1 CD4^+^ T cells and cytotoxic CD8^+^ T cells (7, 16-18) while also promoting CD4^+^ and CD8^+^ Treg responses (16, 18-20). These properties have enabled treatments with GABA-R agonists to inhibit the progression of a diverse array of autoimmune diseases that occur in mice with different genetic backgrounds, including models of type 1 diabetes (T1D), multiple sclerosis (MS), rheumatoid arthritis, Sjogren’s syndrome, as well as inflammation in type 2 diabetes (9, 13, 17, 18, 20-22). Relevant to the treatment of COVID-19, GABA inhibits human immune cell secretion of many pro-inflammatory cytokines and chemokines associated with cytokine storms in individuals with COVID-19 (5, 7, 23-27).

Airway epithelial cells and type II alveolar epithelial cells also express GABA_A_-Rs which modulate the intracellular ionic milieu and help maintain alveolar fluid homeostasis (28-31). GABA and GABA_A_-R positive allosteric modulators (PAMs) reduce inflammation and improve alveolar fluid clearance and lung functional recovery in different rodent models of acute lung injury (32-39), and limit pulmonary inflammatory responses and improve clinical outcomes in ventilated human patients (40-42). GABA reduces human bronchial epithelial cell secretion of inflammatory factors *in vitro* (33) and GABA_A_-R PAMs reduce macrophage infiltrates and inflammatory cytokine levels in bronchoalveolar lavage fluid and reduce rodent and human macrophage inflammatory responses (8, 11, 29, 38, 43-45). Moreover, GABA inhibits human neutrophil activation (46) and platelet aggregation (47), which is potentially important because pulmonary thrombosis often occurs in critically ill COVID-19 patients. Finally, lower levels of plasma GABA are detected in hospitalized COVID-19 patients and associated with the pathogenesis of COVID-19 (48), although the basis for this observation is not understood. Together, the multiple actions of GABA_A-_R modulators on various immune cells and lung epithelial cells, along with their safety for clinical use, make them candidates for limiting dysregulated immune responses, severe pneumonia, and lung damage due to coronavirus infection.

In a previous study, we began to assess whether GABA-R agonists had therapeutic potential for treating coronavirus infections by studying A/J mice that were inoculated with mouse hepatitis virus (MHV-1) (49). MHV-1 is a pneumotropic beta-coronavirus that induces a highly lethal pneumonitis in A/J mice (50-53). We observed that oral administration of GABA or the GABA_A-_R-specific agonist homotaurine, but not a GABA_B-_R-specific agonist, effectively inhibited MHV-1-induced pneumonitis, disease severity, and death rate when given before or after the onset of symptoms (49). GABA treatment also reduced MHV-1 viral load in their lungs.

Based on the above findings, we sought to test the therapeutic potential of GABA treatment in a model that was more clinically relevant to COVID-19. Accordingly, we assessed the ability of GABA treatment to limit the disease process in SARS-CoV-2 infected transgenic mice expressing human ACE2 (K18-hACE2 mice) which provides an acute and lethal model of COVID-19 (54-59).

## Results

### GABA treatment protects SARS-CoV-2-infected K18-hACE2 transgenic mice from severe illness and death

K18-ACE2 mice were inoculated with SARS-CoV-2 intranasally and placed in cages with water bottles containing 0, 0.2, or 2.0 mg/mL GABA, as described in Methods. The mice were maintained on these treatments for the entire study. We monitored the animal’s behavior and scored the severity of the disease twice daily after infection. There was no statistically significant difference in longitudinal body weights or the amounts of water consumed among the different groups of mice (Supplementary Fig. 1). The SARS-CoV-2 infected control mice that received plain water developed overt signs of illness beginning around day 5 post-infection which increased in severity and often necessitated humane euthanasia prior to the end of the study (Fig. 1A, 1B), similar to previous observations in this model (57). In contrast, the mice receiving GABA displayed significantly reduced illness scores compared to the control mice (Fig. 1A). Only 20% of control mice survived until the pre-determined end of the studies 7-8 days post-infection. In contrast, 80% of the mice that received GABA at 0.2 or 2 mg/mL were surviving on days 7-8 post-infection (Fig. 1B, p=0.004 and p=0.018, respectively vs. the control).

**Fig. 1.**
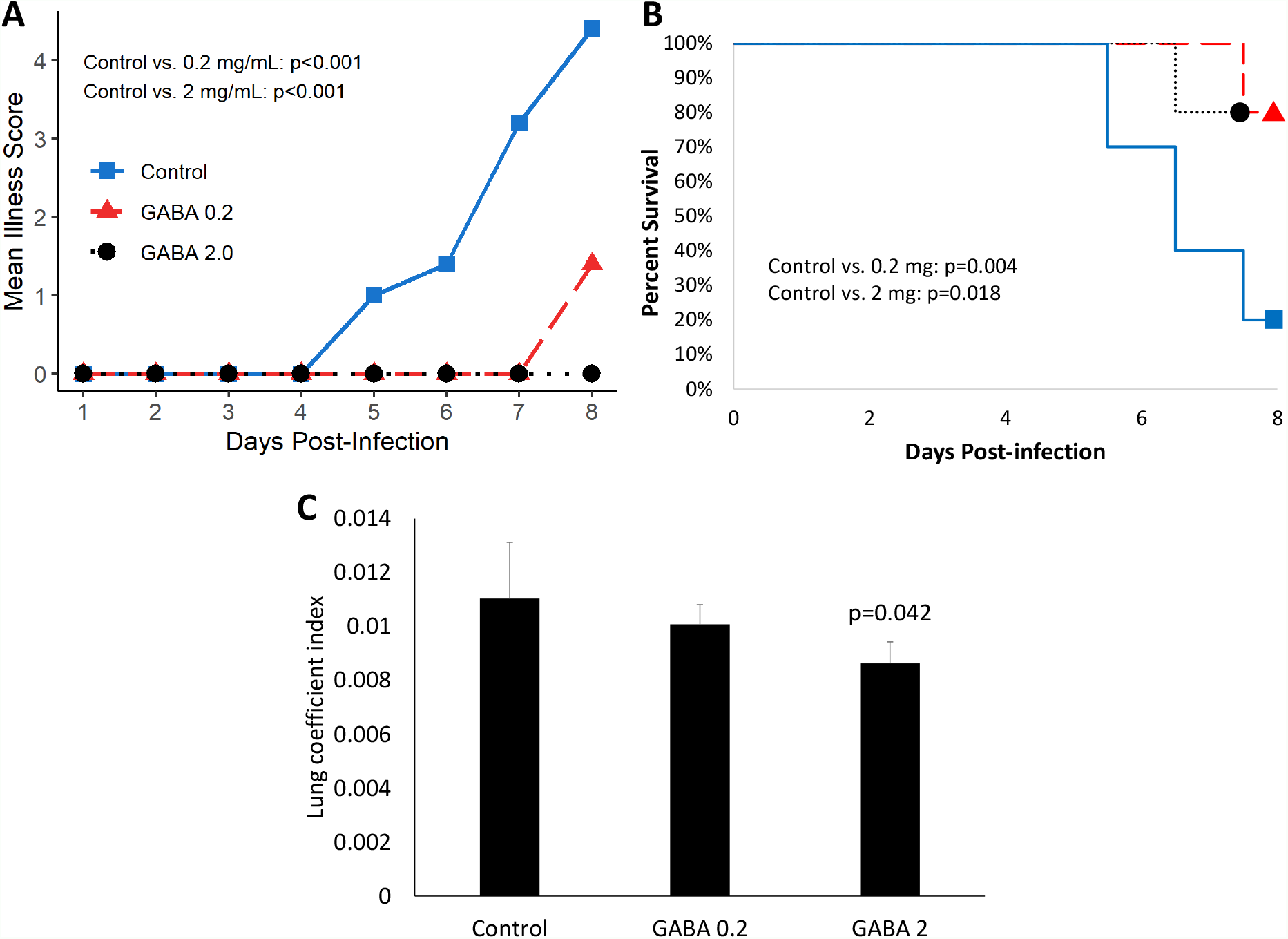
GABA treatment limits disease severity and mortality in SARS-CoV-2 infected K18-hACE2 mice. Following SARS-CoV-2 inoculation the mice were placed on plain water (blue line with squares), 0.2 mg/mL GABA (red line with triangles), or 2.0 mg/mL GABA (black dotted line with circles). **A**. Longitudinal mean illness scores. Disease severity was scored in one of the studies and compared between groups by fitting mixed-effect linear regression models with group and time as fixed effects (to compare means), and with group, time and group by time interaction as fixed effects (to compare slopes). Mouse ID was used as a random effect. N=5 mice/group. **B**. Combined percent surviving mice from two independent studies with 5 mice/group which followed the mice for 7 or 8 days post-infection (n=10 mice/group total). Survival curves were estimated using the Kaplan-Meier method and statistically analyzed by the log-rank test. **C**. Mice that reached an illness score of 5 or survived to the end of the observation period were euthanized. Their lungs were dissected and weighed to calculate the lung coefficient index (the ratio of lung weight to body weight). The data shown are the mean lung coefficient index ± SEM/ N=5 mice/group. The p-value was determined by Student’s t-test.

At the time of euthanasia, each animal’s lungs were excised and weighed. The ratio of lung weight to total body weight (i.e., the lung coefficient index) reflects the extent of edema and inflammation in the lung. We observed a significant reduction in the lung coefficient index scores in the group treated with 2.0 mg/mL GABA (Fig. 1C), indicating reduced lung inflammation. Together, these studies demonstrate that GABA treatment reduces disease severity, inflammation, and death rates in SARS-CoV-2 infected mice.

### GABA treatment reduces SARS-CoV-2 titers in the lungs of infected K18-hACE2 mice

We next asked whether GABA’s protective mechanism could be due in part to reducing viral replication in their lungs. K18-hACE2 mice were infected with SARS-CoV-2 (2,000 TCID_50_ intranasally) and placed on plain water (control) or water containing GABA (2 mg/mL) for the rest of the study. Three days post-infection, which should be near to the peak of viral load in the lungs (57), we harvested their lung tissues to determine viral load by TCID_50_ assay using Vero E6 cells. We found that SARS-CoV-2 titers in the lungs of GABA-treated mice were on average a 1.36 log10 (23-fold) lower than that in the lungs of mice that received plain water (p<0.0001, Fig. 2). Thus, GABA treatment reduced viral loads in the lungs of SARS-CoV-2 infected mice.

**Fig. 2.**
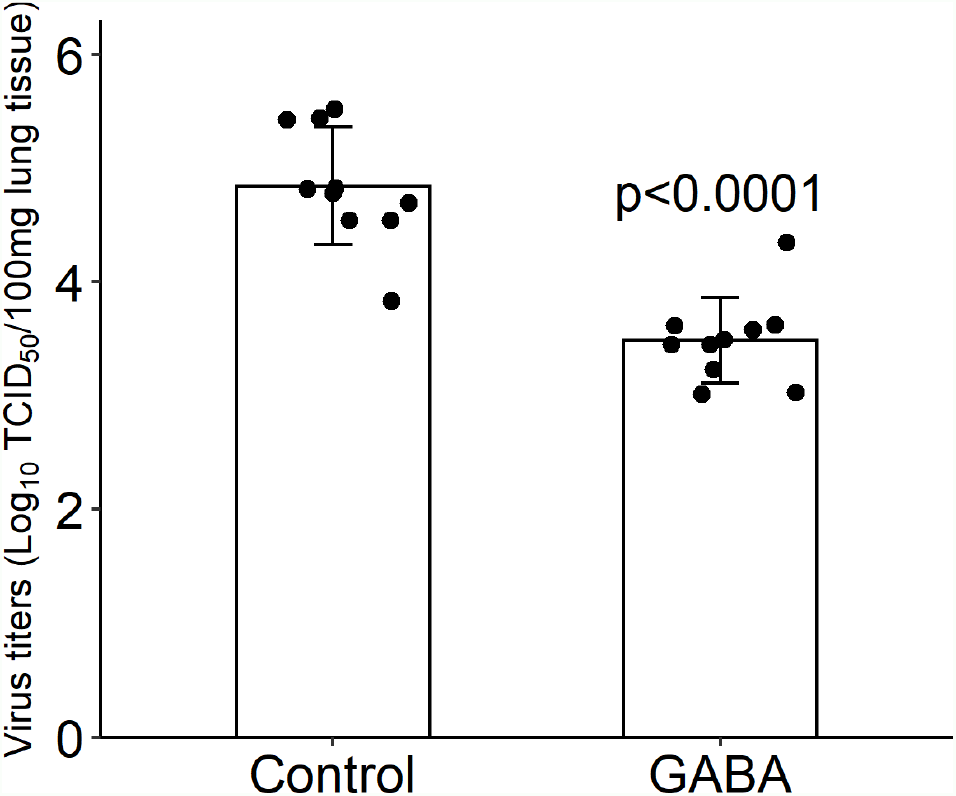
GABA treatment reduces SARS-CoV-2 titers in the lungs. Three days post-infection the right lung was harvested from each mouse and homogenized. The viral titer in lung tissue (100 mg) were determined by TCID_50_ assay using Vero E6 cells. Black dots show viral titers for individual mice (determined in quadruplicate). The data shown are the mean virus titer (Log_10_ TCID_50_/100 mg lung tissue) ± SD in mice given plain water (control) or GABA. N=10 mice/group. The p-value was determined by Student’s t-test.

### GABA treatment modulates early cytokine and chemokine responses to SARS-CoV-2 infection

We next asked whether GABA treatment could modulate serum cytokine and chemokine levels in SARS-CoV-2 infected mice. Sera samples were collected three days post-infection from the same mice that were used to study lung viral load and were analyzed using a multiplex bead kit designed to detect 13 cytokines and chemokines. We did not detect any statistically significant difference in the levels of serum IFNα, IFNβ, IFNγ, IL-12, or GM-CSF between SARS-CoV-2 infected mice that did or did not receive GABA treatment (Fig. 3). There was, however, some suggestion that GABA treatment elevated type I interferons in some mice because only 1/10 mice in the healthy control (HC) and in the SARS-CoV-2-infected control (SC) group had a detectable level of IFNß, while 3/10 of the infected mice which received GABA (G) had detectable IFNß levels and these were greater in magnitude than that found in the other groups (Fig. 3). The median and the mean levels of IFNα were also slightly elevated in the GABA treated vs. untreated SARS-CoV-2 infected group (Fig. 3). Conversely, 3/10 mice in the SC group displayed high levels of IFNγ vs 1/10 mice in the G group and the median and mean IFNγ levels were reduced in the GABA-treated vs. untreated SARS-CoV-2 infected mice (Fig. 3, p=0.25)

**Fig. 3.**
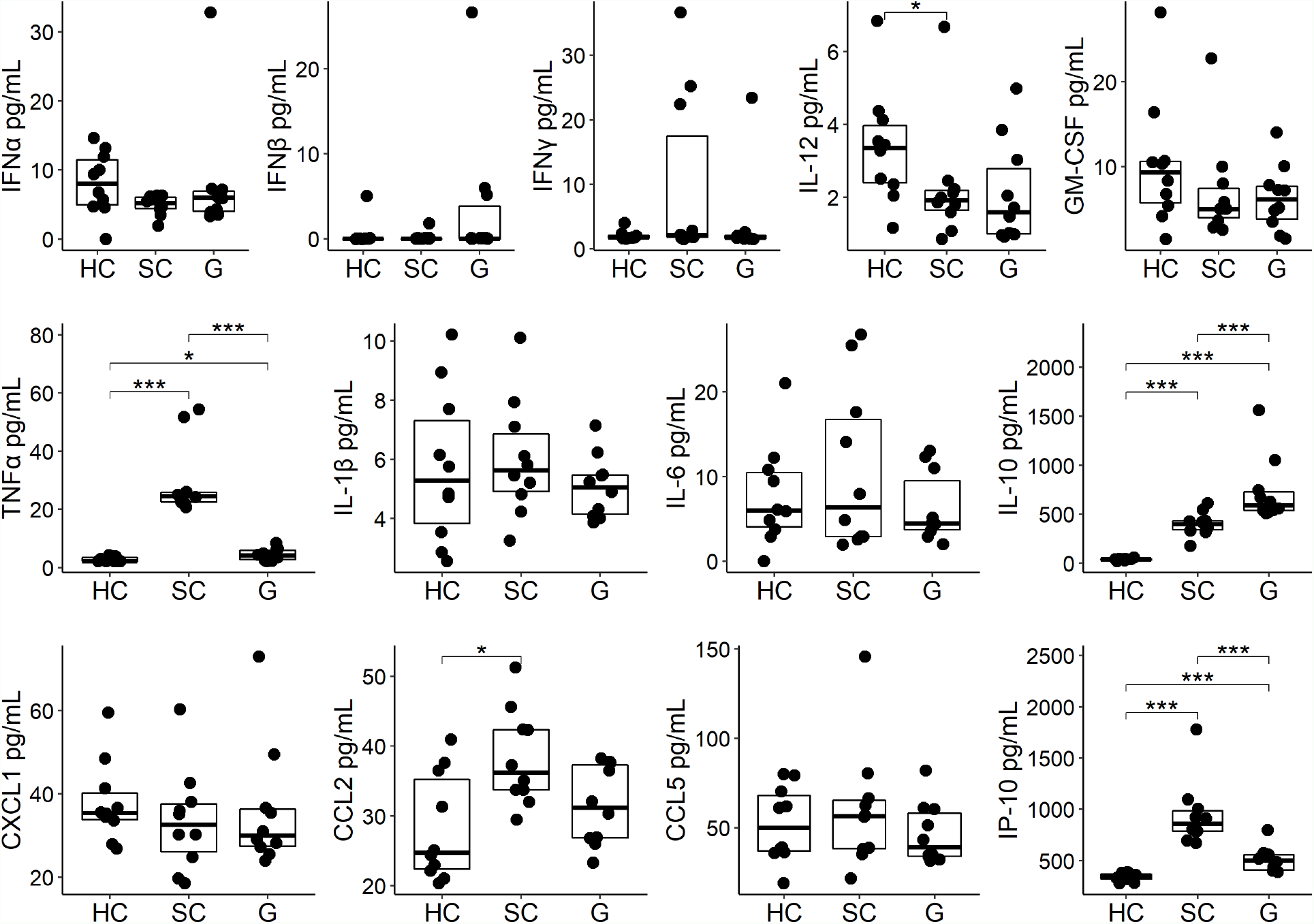
GABA treatment modulates circulating cytokines and chemokines in SARS-CoV-2-infected K18-hAC2 mice. Three days post-infection sera from individual SARS-CoV-2 infected control (SC) mice that were untreated, or were GABA treated (G), were collected and frozen at -80 °C until analysis by a multiplex assay as described in Methods. In addition, sera from age-matched health control (HC) B6 mice were studied. The non-normally distributed data are shown in boxplots with the borders of the box indicating 1st and 3rd quartile of each group (n=10), the bolded line indicating the median. The data were analyzed by Wilcoxon rank-sum tests. *p<0.05, ***p<0.001

Analysis of pro-inflammatory and anti-inflammatory cytokines revealed that SARS-CoV-2 infection significantly increased the levels of serum TNFα in the SC group (p<0.001, Fig. 3) as in previous studies of SARS-CoV-2-infected K18-hACE2 mice (57, 59-61). GABA treatment significantly mitigated TNFα production following infection (p<0.001). There was no statistically significant difference in the levels of serum IL-1β and IL-6 although the levels of serum IL-1β and IL-6 in the GABA-treated group were slightly reduced compared with the SC group (Fig. 3). SARS-CoV-2 infection significantly increased the levels of serum IL-10, as in past studies of K-18-hACE2 mice (57, 59, 61). GABA treatment further significantly elevated the serum IL-10 levels above that in untreated SARS-CoV-2 infected mice (p<0.001).

Analysis of serum chemokines revealed that untreated SARS-CoV-2 infected mice, but not the GABA-treated SARS-CoV-2 infected mice, displayed significantly higher levels of CCL2 relative to healthy controls (p<0.05), with the levels of CCL2 tending to be lower in GABA-treated vs. untreated-infected mice (p=0.07). Moreover, GABA treatment led to significantly reduced levels of IP-10 (CXCL10) compared to that in untreated SARS-CoV-2 infected mice (p<0.001, Fig. 3). We observed no statistically significant difference in the levels of serum CXCL1 and CCL5 among these groups of mice. Thus, GABA treatment shifted systemic cytokine and chemokine responses towards those associated with less risk for developing severe COVID-19 by increasing early type I IFN responses in some mice, significantly reducing the levels of TNFα and IP-10, tending to reduce CCL2 levels, but enhancing IL-10 production in SARS-CoV-2 infected mice

### GABA treatment protects SARS-CoV-2-infected mice from severe illness when treatment is initiated near the peak of viral load in the lungs

We next examined whether GABA treatment could be also beneficial when treatment was initiated at a later stage of the disease process. K18-hACE2 mice were infected with SARS-CoV-2 as above and GABA treatment (2 mg/mL) was initiated at 2 days post-infection, near the peak of viral load in the lungs. Compared to untreated control mice, GABA-treated mice displayed reduced longitudinal mean illness scores (overall p=0.002, Fig. 4A). At the end of the study on day 7, only 22% of control mice survived, while 88% of GABA-treated mice had survived (p=0.004, Fig. 4B). GABA-treated mice also had a reduced lung coefficient index (p<0.001, Fig. 4C).

**Fig. 4.**
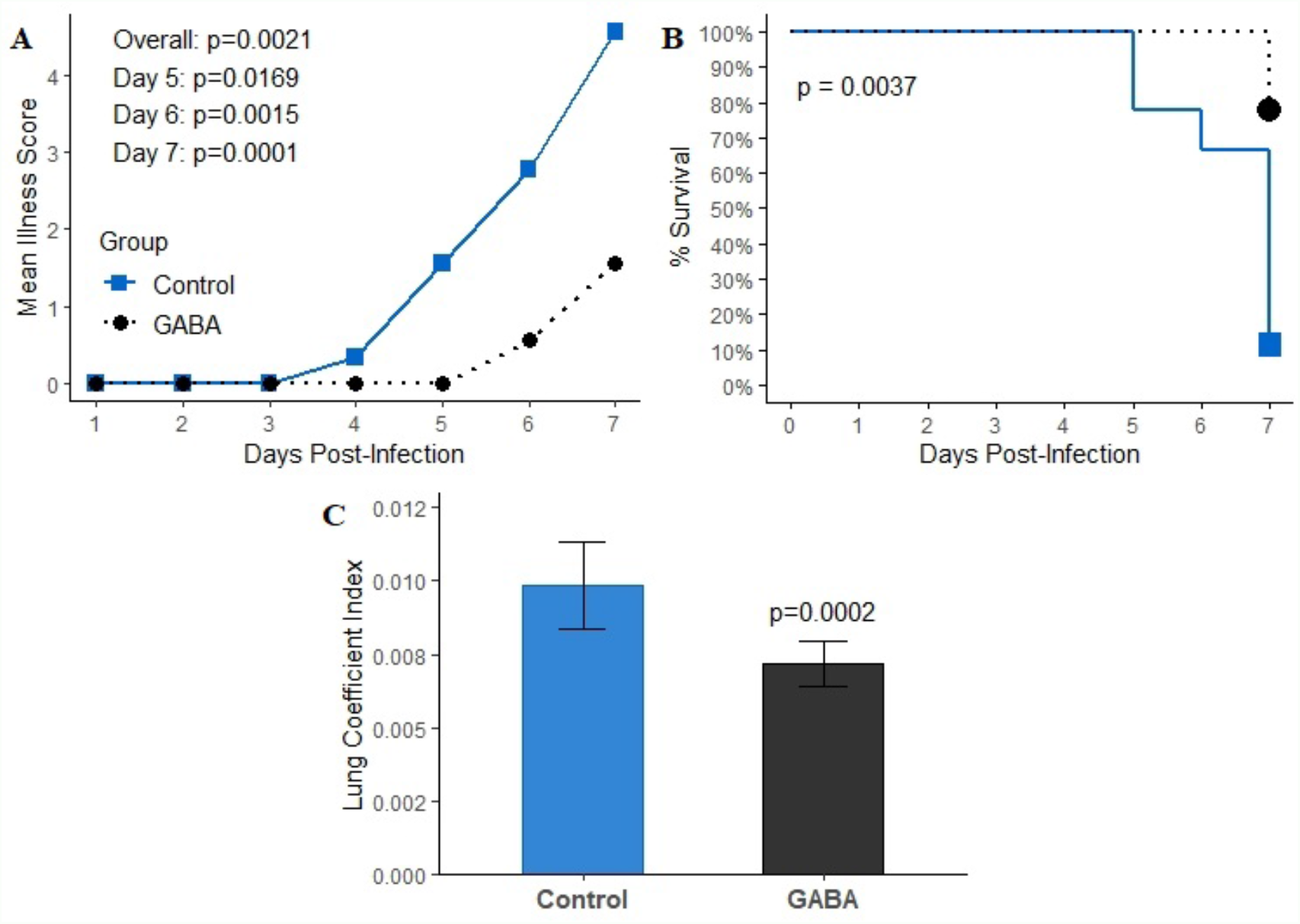
GABA treatment initiated near the peak of lung viral load has protective effects in SARS-CoV-2 infected K18-hACE2 mice. Mice were infected with SARS-CoV-2 and two days later the mice received GABA through their drinking water (2 mg/mL), or were maintained on plain water for the remainder of the study. **A**. Longitudinal mean illness scores. The data were analyzed by fitting mixed-effect linear regression models with group and time as fixed effects (to compare means), and with group, time and group by time interaction as fixed effects (to compare slopes). Mouse ID was used as a random effect. **B**. Percent surviving mice. The data were analyzed by Kaplan-Meier survival curves and the log-rank test. **C**. Lung coefficient index. The data were analyzed by Student t-test. N=9 mice/group.

## Discussion

Our studies demonstrated that GABA administration initiated immediately, or two-day post-SARS-CoV-2 infection, reduced the lung coefficient index, lung viral load, and death rate in SARS-CoV-2 infected mice. These results, along with our previous findings in the MHV-1 infected mice, are the first reports of GABA administration modulating the outcome of a viral infection. GABA’s beneficial effects in these two different highly lethal coronavirus models suggest that GABA-R activation may be a generalizable therapeutic strategy to help reduce the severity of coronavirus pulmonary infections, at least in mice.

Our observations are surprising in a number of ways. First, based on GABA’s anti-inflammatory actions in models of autoimmune disease, cancer, and parasitic infection, it was highly possible that early GABA treatment could have exacerbated the disease in SARS-CoV-2 infected mice by limiting or delaying the innate immune responses to the viral infection. However, SARS-CoV-2-infected mice that received GABA treatment at the time of infection, or 2-days post-infection, fared much better than their untreated SARS-CoV-2-infected counterparts.

Second, GABA’s ability to modestly reduce viral load in the lungs was surprising since the traditional targets of antiviral medications are viral proteases or polymerases. We know from large-scale screens of drug libraries that GABA-R agonists do not interfere with SARS-CoV-2 binding to ACE2 or its internalization into cultured Vero E6 cells (62). Notably, GABA_A_-Rs are expressed by lung bronchial and alveolar cells (28, 29, 63) and it is possible that GABA_A_-R activation led to changes in intracellular ion levels that made the environment less favorable for viral replication. While the activation of neuronal GABA_A_-Rs leads to Cl^-^ influx and hyperpolarization, the activation of GABA_A_-Rs on other types of cells, such as alveolar ATII cells, causes Cl^-^ efflux and depolarization (28, 29). Many viruses, including some coronaviruses, can promote Ca^2+^ influx into their host cell to enhance their replication (64, 65). The activation of GABA_A_-Rs on infected cells could promote Cl^-^ efflux which would oppose Ca^2+^ influx and reduce Ca^2+^ contents, limiting SARS-CoV-2 replication. Indeed, calcium blockers have been shown to reduce SARS-CoV-2 replication *in vitro*, but whether they confer beneficial effects to COVID-19 patients has been controversial (66-68). The activation of GABA_A_-Rs on alveolar and large airway epithelial cells may also have altered 1) the secretion of inflammatory signaling molecules from infected cells, 2) alveolar surfactant production/absorption, and/or 3) altered inflammatory responses and autophagy processes (15) in ways that limited virus infection and replication.

Third, GABA treatment shifted some cytokine and chemokine levels in directions that are expected to be beneficial if extended to COVID-19 treatment. First, early GABA treatment elevated type 1 interferons in some mice. Since delayed or reduced type I interferon responses are a risk factor for developing severe COVID-19 (69), such tendencies could be beneficial.

GABA treatment significantly reduced circulating TNFα levels in infected mice, extending previous observations that GABA inhibits the NF-κB activation in mouse and human immune cells (7, 70). As a result, GABA treatment slightly decreased mean serum IL-6 from 10.7 to 6.3 pg/mL. Since TNFα and IL-6 are important pro-inflammatory mediators, the decreased levels of circulating TNFα and IL-6 indicated that GABA treatment suppressed innate immune responses, which is likely to have contributed to its protective effects.

GABA-treated mice also had reduced levels of serum IP-10, a pro-inflammatory chemokine that attracts the migration of CXCR3^+^ macrophages/monocytes, T cells, and NK cells (71). Elevated levels of IP-10 are consistently detected in severely ill COVID-19 patients and may provide a predictive marker of patient outcome (72-75). IP-10 production is induced by IFNγ, NF-κB activation and other stimulators in several types of cells (76). Consistent with the reduced IP-10 levels, we also found that the serum mean IFNγ level in the GABA-treated mice was about half that in the untreated SARS-CoV-2 infected mice (4.0 vs. 9.8 pg/mL). These data suggest that early GABA treatment reduced IFNγ production and together with its inhibition of NF-κB activation, led to decreased IP-10 secretion. Given that IP-10 functions to recruit inflammatory cell infiltration into lesions and modulates cell survival, the lower levels of circulating IP-10 in GABA-treated mice are likely to have limited the migration of macrophages, monocytes, and NK cells into the pulmonary lesions and helped to protect the mice from death. Similarly, we also observed that GABA treatment slightly reduced the levels of serum CCL2 which may have contributed to protecting mice from death since high levels of CCL2 are associated with high mortality in COVID-19 patients (77).

GABA treatment also enhanced IL-10 levels in SARS-CoV-2 infected mice. IL-10 is generally regarded as an anti-inflammatory cytokine, however, it can be immunostimulatory in certain contexts and elevated levels of IL-10 are associated with the development of severe COVID-19 (78, 79). If the elevation of IL-10 levels in GABA-treated SARS-CoV-2 infected mice had counter-therapeutic effects, it is evident that GABA’s beneficial actions were functionally dominant leading to improved outcomes. Conceivably, the enhanced levels of IL-10 levels due to GABA treatment may have been therapeutic by 1) its classical anti-inflammatory actions, 2) exhausting immune cells, 3) reducing tissue damage in the lungs, or 4) other yet to be identified actions.

Initiating GABA treatment 2-days post-infection, near the peak of viral loads in the lungs, also decreased disease severity and death rates during the observation period. This observation bodes well for the clinical potential of this therapeutic strategy. It will be of interest to study the impact of GABA treatment on inflammation in the lungs when treatment is initiated at even later time points post-infection.

Besides expressing the hACE2 transgene in lung cell epithelial cells, K18-hACE2 mice express hACE2 ectopically in their CNS leading to the spreading of SARS-CoV-2 infection to their CNS at late stages of the disease process (57, 59, 60). Because GABA does not pass through the blood-brain barrier (BBB), it is unlikely that GABA treatment directly exerted beneficial effects in the CNS. However, the decreased levels of circulating proinflammatory cytokines and chemokines in GABA-treated mice may have also reduced their entry into the CNS. While SARS-CoV-2 is thought to not efficiently replicate in the human CNS (80), some COVID-19 patients experience cognitive impairments (or “brain fog”). Histological studies of the brains from COVID-19 patients have observed immune cell infiltrates and increased frequencies of glial cells with inflammatory phenotypes which are indicative of neuroinflammatory responses (80-87). In previous studies, we have shown that treatment with homotaurine, a GABA_A_-R-specific agonist that can pass through the BBB, reduced the spreading of inflammatory T cell responses within the CNS, limited the pro-inflammatory activity of antigen-presenting cells, and ameliorated disease in mouse models of multiple sclerosis (13, 18). Homotaurine may have also down-regulated the inflammatory activities of GABA_A_-R-expressing microglia and astrocytes in the CNS (88). Homotaurine was as effective as GABA in protecting MHV-1-infected mice from severe illness, pointing to GABA_A_-Rs as the major mediators of GABA’s beneficial effects in this model (49). These observations suggest that homotaurine treatment may provide a new strategy to reduce inflammation in the CNS stemming from COVID-19 or inflammatory neurological disorders. Homotaurine (also known as Tramiprosate) was found to physically interfere with amyloid aggregation *in vitro*, leading to its testing as a treatment for Alzheimer’s disease in a large long-term phase III clinical study (89-91). While this treatment failed to meet primary endpoints, the treatment appeared to be very safe and follow-up studies suggested some disease-modifying effect (92).

Interestingly, circulating GABA levels are significantly reduced in hospitalized patients with COVID-19 (48). This clinical finding, independent of our results presented here, raises the question of whether GABA therapy could be beneficial for COVID-19 patients. Many individuals take GABA_A_-R PAMs (e.g., Xanax), however, these PAMs are not expected to be useful for COVID-19 treatments because they only extend the opening of the GABA_A-_R Cl^-^ channel after it is opened by GABA. GABA levels in the blood appear to be too low to provide protection in models of autoimmune disease and coronavirus (MHV-1 and SARS-CoV-2) infection as evidenced by the need to administer GABA exogenously for a therapeutic effect in these models.

It is likely that SARS-CoV-2 variants and novel coronaviruses will constantly arise that will be insufficiently controlled by available vaccines and antiviral medications. Developing new vaccines against these new viruses will be much slower than the spread of these new viruses among the world population. Our findings suggest that GABA-R agonists may provide inexpensive off-the-shelf agents to help lessen the severity of disease caused by these new viruses. Because GABA’s mechanisms of action are unlike that of other coronavirus treatments, combination treatments could have enhanced benefits.

GABA is regarded as safe for human use and is available as a dietary supplement in the USA, China, Japan, and much of Europe (93). In other countries, because GABA is a non-protein amino acid, it is regulated as a medicinal agent or drug (e.g., in the UK, Canada, and Australia). In our studies, GABA at 2.0 and 0.2 mg/mL were equally effective at protecting SARS-CoV-2 infected mice from death (Fig. 1B). The human equivalent dose of GABA at 0.2 mg/mL (assuming consumption of 3.5 mL/day water, see Supplemental Fig. 1 and calculated as per (94)) is 0.68 g/day for a 70 kg person, which is well within the level known to be safe (93). While our preclinical observations indicate that GABA-R agonists are promising candidates to help treat coronavirus infections, information on their dosing and the time window during which their effects might be beneficial or deleterious during a coronavirus infection in humans are lacking and clinical trials are needed to assess their therapeutic potential.

## Limitations of this study

There are a number of major limitations of this study. First, the K18-hACE2 mouse model imperfectly models SARS-CoV-2 infection and immune responses in humans. Second, GABA’s impact on immune cells and lung cells may differ in important ways between mice and humans. Third, GABA treatment may only be beneficial during a specific time window of the disease process and at other times may be deleterious. Accordingly, careful clinical trials are needed to determine the time window and dosage, if any, that GABA-R agonist treatment has a beneficial effect in COVID-19 patients.

## Materials and Methods

### Virus

SARS-CoV-2 (USA-WA1/2020) was obtained from the Biodefense and Emerging Infections Resources of the National Institute of Allergy and Infectious Diseases. All *in vivo* studies of SARS-CoV-2 infection were conducted within a Biosafety Level 3 facility at UCLA or USC. SARS-CoV-2 stocks were generated by infection of Vero-E6 cells (American Type Culture Collection (ATCC CRL1586)) cultured in DMEM growth media containing 10% fetal bovine serum, 2 mM L-glutamine, penicillin (100 units/ml), streptomycin (100 units/ml), and 10 mM HEPES. The cells were incubated at 37°C with 5% CO_2_ and virus titers were determined as described below for tittering virus in lung homogenates.

### Mice and GABA treatments

Female K18-hACE2 mice (8 weeks in age) were purchased from the Jackson Laboratory. The first survival study was conducted within USC’s BSL3 Core facility and all following studies were conducted in the UCLA ABSL3 Core Facility. One week after arrival, they were inoculated with SARS-Cov-2 (2,000 PFU or 2,000 TCID_50_ (as per (57) at USC and UCLA, respectively) in 20 µl Dulbecco’s modified Eagle’s medium. The mice were randomized into groups of 5-9 mice and were placed on water bottles containing 0, 0.2, or 2.0 mg/mL GABA (Millipore-Sigma (stock #A2129), St. Louis, MO, USA) immediately, or 2 days post-infection (as indicated) and maintained on those treatments for the remainder of each study. These studies were carried out in accordance with the recommendations of the Guide for the Care and Use of Laboratory Animals of the National Institutes of Health. The protocols for all experiments using vertebrate animals were approved by the Animal Research Committee at UCLA (Protocol ID: ARC #2020-122; 8/25/20-8/24/2023) or USC (IACUC protocol # 21258; 1/28/2021-1/27/2024) and were carried out in compliance with the ARRIVE guidelines.

### Illness scoring and lung index score

Individual mice were monitored twice daily by two to three observers for their illness development and progression which were scored on the following scale: 0) no symptoms, 1) slightly ruffled fur and altered hind limb posture; 2) ruffled fur and mildly labored breathing; 3) ruffled fur, inactive, moderately labored breathing; 4) ruffled fur, inactive, obviously labored breathing, hunched posture; 5) moribund or dead.

The percent survival of each group of mice was determined longitudinally. Mice with a disease score of 5 were weighed, euthanized, and their lungs removed and weighed for calculation of lung coefficient index (the ratio of lung weight to total body weight, which reflects the extent of edema and inflammation in the lungs). On day 7 or 8 post-infection, the surviving animals were weighed, euthanized, and their lungs were removed and weighed for determination of the lung coefficient index.

### Viral titers in lungs

Female K18-ACE2 mice (9 weeks in age) were intranasally inoculated with 2,000 TCID_50_ SARS-Cov-2 and placed on plain water or water containing GABA (2 mg/mL) for the rest of the study. Three days post-infection, the mice were euthanized and individual blood samples were collected for preparing serum samples. A portion of their lower right lung was weighed and about 100 mg of wet lung tissue from each mouse was homogenized into 1 mL of ice-cold DMEM with 10% fetal calf serum (FBS) with 1 mm glass beads using a Minilys homogenizer at 50 Hz for 90 seconds, followed by centrifugation. Their supernatants were collected for virus tittering. Vero E6 cells (1×10^4^ cells/well) were cultured in DMEM medium supplemented with 10% FBS in 96-well plates overnight to reach 80% of confluency and infected in quintuplicate with a series of diluted mouse lung homogenates in 100 µl of FBS-free DMEM medium at 37º C for four days. The percentages of viral cytopathic effect areas were determined. The SARS-CoV-2 titers were calculated by the Reed and Muench method (95).

### Simultaneous detection of multiple cytokines and chemokines by flow cytometry

Blood samples were collected from the same mice that were used to study viral loads in the lungs at three days post-infection. Sera from individual mice were prepared and stored at -80 °C. The levels of serum cytokines and chemokines were determined by a bead-based multiplex assay using the LEGENDplex mouse anti-virus response panel (13-plex) kit (#740622, Biolegend, San Diego, USA), according to the manufacturer’s instructions. Briefly, the control and experimental groups of serum samples were diluted at 1:2 and tested in duplicate simultaneously. After being washed, the fluorescent signals in each well were analyzed by flow cytometry in an ATTUNE NxT flow cytometer (Thermofisher). The data were analyzed using the LEGENDplex™ Data Analysis Software Suite (Biolegend) and the levels of each cytokine or chemokine in serum samples were calculated, according to the standard curves established using the standards provided.

### Statistics

Statistical methods are described in each figure legend. A p-value of <0.05 was considered statistically significant.

## Acknowledgments

We would like to thank Dr. Vaithilingaraja Arumugaswami for his kind provision of virus stock and Drs. Jennifer Hahn, Orian Shirihai, Anton Petcherski, Theodoros Kelesidis, and Joanne L. Zahorsky-Reeves, as well as Madhav Sharma, Melanie Ciampaglia, Tomoko Yamada, and Lenore Kaufman for their help. We thank Lucia Chen, Dr. Jeffery Gornbein, and Dr. Alexandra Klomhaus for statistical analysis and figure preparation, and Miss. Salem Haile in the Janice Gorgi Flow Cytometry Center for FACS analysis. We thank Jeremy Marshall and Philip Sell for their support in the ABSL3 lab at USC.

## Author Contributions

Conceived and designed the experiments: JT, DLK; Performed the experiments: JT, BD, and JH; Supervised work and assessed data quality in the ABSL3 labs BD, LC. Analyzed the data: JT, DLK. Wrote the paper: JT, DLK. Daniel Kaufman and Jide Tian are guarantors of this work and, as such, had full access to all the data in the study and take responsibility for the integrity of the data and the accuracy of the data analysis. All authors approved the final manuscript as submitted.

## Institutional Review Board Statement

This study was carried out in accordance with the recommendations of the Guide for the Care and Use of Laboratory Animals of the National Institutes of Health. The protocols for all experiments using vertebrate animals were approved by the Animal Research Committee at UCLA (Protocol ID: ARC #2020-122; Date 8/25/20-8/24/2023) and USC (IACUC protocol # 21258 (1/28/2021-1/27/2024), IBC: BUA 20-0030/ 20-0033)

## Data Availability Statement

The data presented in this study are available on request from the corresponding authors.

## Supplemental data

**Supplementary Figure 1.**
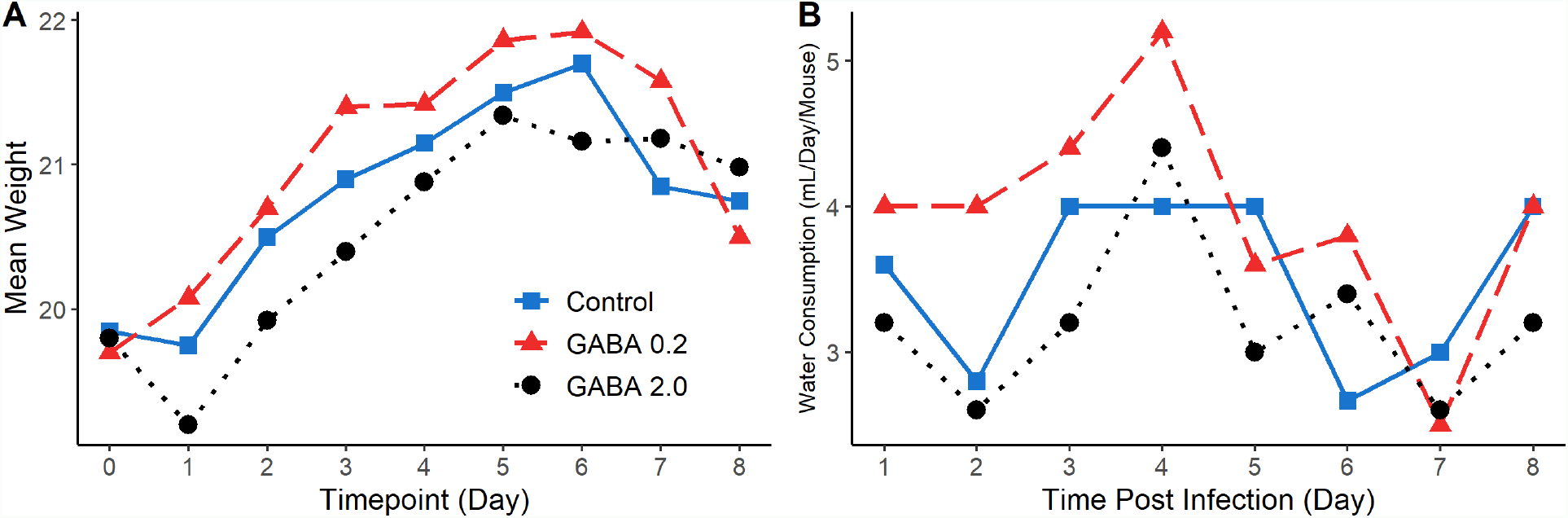
A) Longitudinal body weights of treatment groups following SARS-CoV-2 infection. Following SARS-CoV-2 infection, the mice were placed on plain water or water containing GABA at the indicated dose for the entire observation period. Their body weights were measured daily. Mice were euthanized when they had an illness score of 5 or at the end of the observation period (8 days post-infection). Data shown is mean body weight for surviving mice in each group. **B) Average daily amount of water consumed**. We monitored how much water was consumed daily per cage and divided that by the number of surviving mice in the cage to calculate the average daily water consumption per mouse. For each treatment, we began with 5 mice/group with each group was housed in a single cage. Data shown is the average daily water consumption (mL) per mouse post-infection.

